# Genetic basis of reproductive isolation in Torrey pine (*Pinus torreyana* Parry): insights from hybridization and adaptation

**DOI:** 10.1101/2024.04.15.589509

**Authors:** Lionel N. Di Santo, Alayna Mead, Jessica W. Wright, Jill A. Hamilton

**Author notes:** Contributed equally to the study (Lionel N. Di Santo, Alayna Mead).

## Abstract

Tree species are often locally adapted to their environments, but the extent to which environmental adaptation contributes to incipient speciation is unclear. One of the rarest pines in the world, Torrey Pine (*Pinus torreyana* Parry), persists naturally across one island and one mainland population in southern California. The two populations are morphologically and genetically differentiated, but experience some connectivity, making it an ideal system for assessing the evolution of reproductive isolation. Previous work has found evidence of heterosis in F1 mainland-island hybrids, suggesting genetic rescue could be beneficial in the absence of reproductive barriers. Using ddRADseq and GWAS for a common garden experiment of island, mainland, and F1 individuals, we identified candidate loci for environmentally-driven reproductive isolation, their function, and their relationship to fitness proxies. By simulating neutral evolution and admixture between the two populations, we identified loci that exhibited reduced heterozygosity in the F1s, evidence of selection against admixture. SNPs with reduced F1 heterozygosity were enriched for growth and pollination functions, suggesting genetic variants that could be involved in the evolution of reproductive barriers between populations. One locus with reduced F1 heterozygosity exhibited strong associations with growth and reproductive fitness proxies in the common garden, with the mainland allele conferring increased fitness. If this locus experiences divergent selection in the two natural populations, it could promote their reproductive isolation. Finally, although hybridization largely reduced allele fixation in the F1s initially, indicating heterosis is likely due to the masking of deleterious alleles, the emergence of reproductive isolation between populations may diminish the longer-term benefits of genetic rescue in F2 or advanced-generation hybrids. As Torrey pine is a candidate for interpopulation genetic rescue, caution is warranted where longer-term gene flow between diverged populations may result in reduced fitness if barriers have evolved.

## Background

Forest trees are well-known for maintaining signatures of local adaptation among populations even when experiencing high levels of homogenizing gene flow (Petit and Hampe 2006; Janes and Hamilton 2017; Cannon and Petit 2020). This apparent paradox is explained by strong competition and selection at early life stages: deleterious non-adaptive alleles introduced by migrant individuals or pollen are likely to be removed from the gene pool (Petit and Hampe 2006). In some cases, population differentiation may lead to reproductive isolation and eventually speciation (Schluter 2001; Hendry et al. 2007; Schluter 2009; Hendry 2009; Nosil et al. 2009b; Andrew and Rieseberg 2013). However, the evolutionary and genomic mechanisms underlying the shift from local adaptation with gene flow to the evolution of reproductive isolation is poorly documented in trees, particularly conifers (Bolte and Eckert 2020). The prevalence of local adaptation even under high levels of gene flow makes trees a good system for better understanding how and when divergent selection may drive the evolution of reproductive isolation, a prerequisite for speciation.

Speciation requires the evolution of genetic divergence, which can arise in response to environmental differences, as a result of isolation, or a combination of both (Schluter 2001; Nosil et al. 2009b,a). Allopatry may cause intrinsic reproductive incompatibilities to evolve between populations, resulting in reproductive isolation if they come into secondary contact. However, reproductive isolation can evolve even when gene flow is occurring if selection against migrants and hybrids is strong enough (Schluter 2009). If non-local alleles are deleterious, selection against them may favor the evolution of reproductive barriers. Extrinsic postzygotic reproductive barriers may occur when environmental selection results in lower hybrid or migrant fitness (as found by Lowry, Rockwood, et al., 2008; Melo et al., 2014; Richards & Ortiz-Barrientos, 2016), while intrinsic postzygotic barriers result in lower hybrid fitness regardless of the environment (Coughlan and Matute 2020). While both types of barriers can be involved in speciation, with great variation in the strength of each type of barrier across systems, extrinsic postzygotic barriers are generally stronger than intrinsic postzygotic barriers (Christie et al. 2022). Additionally, extrinsic postzygotic barriers are stronger in ecotypes than in species, suggesting they may be more important at earlier stages of speciation (Christie et al. 2022) with intrinsic postzygotic isolation being important at later stages, either as a result of divergence or by contributing to reinforcement (Coughlan and Matute 2020).

As reproductive isolation can arise through many mechanisms, including both prezygotic and postzygotic barriers, identifying the genes underlying these reproductive barriers can help us better understand the process of speciation at the genomic level (Lowry et al. 2008a; Rieseberg and Blackman 2010; Feder et al. 2012; Strasburg et al. 2012; Choi et al. 2020; Schluter and Rieseberg 2022). One way to identify mechanisms underlying the evolution of reproductive isolation is through signatures left in the genome (Nosil et al. 2009a; Schluter 2009; Feder et al. 2012; Strasburg et al. 2012). When reproductive isolation is developing amidst ongoing gene flow, as in the early stages of ecological speciation where species diverge as a consequence of natural selection among contrasting environments, loci that are under selection will become more differentiated between two populations than other genomic regions (Feder et al. 2012). As speciation proceeds, variants linked to those regions under selection diverge, eventually leading to differentiation throughout the genome. Analysis of the functions of these divergent regions of the genome can be complemented with phenotypic data where both populations are grown in a common environment. Using genotype-phenotype associations, differentiated loci can be linked to variance in reproductive or fitness-associated traits. As reproductive-aged common gardens are uncommon for long- lived species, our understanding of reproductive barriers and their genomic underpinnings in trees is limited.

In this study, we took advantage of a 10-year-old common garden with Torrey Pine (*Pinus torreyana* Parry), established in 2007 by the USDA-Forest Service Pacific Southwest Research Station. Torrey pine is one of the rarest pines in the world with only two native populations remaining (Di Santo et al. 2022). Separated by approximately 280 km (170 miles), the island population native to Santa Rosa Island, one of the Channel Islands, experiences cooler temperatures and greater precipitation on average than the mainland population located at the Torrey Pine State Natural Reserve in La Jolla, California (e.g., mean temperature of the driest quarter is 16.7 °C at the island population and 18.9 °C at the mainland population; precipitation of the warmest quarter is 17 mm at the island population and 12 mm at the mainland population, Appendix S1 and S2). When grown in a common environment, the two populations exhibit differences in growth, cone, and needle morphology consistent with adaptation to their native environments (Hamilton et al. 2017), but also exhibit relatively low genetic differentiation (F_ST_ = 0.0129) and some evidence of ongoing gene flow (Di Santo et al. 2022). In an open-pollinated common garden including mainland and island trees, the only F1s produced were from island maternal trees fertilized by mainland pollen, with no F1s observed between mainland maternal trees fertilized by island pollen (Hamilton et al. 2017). Asymmetric crossing barriers can be evidence of cytonuclear or maternal incompatibility (Tiffin et al. 2001; Turelli and Moyle 2007; Case et al. 2016), and may be a sign that reproductive barriers between the two populations have already begun to evolve (Barnard-Kubow et al. 2016). Thus, Torrey pine is an ideal system for investigating the early stages of reproductive isolation between two geographically separated populations occurring in contrasting environments. To evaluate the genomic and fitness consequences of inter-population gene flow, we leverage a common garden experiment of reproductive-aged trees of both parental populations and their F1 hybrids.

Understanding the nature of the divergence between Torrey pine populations will be critical to the species’ management. Both populations exhibit exceedingly low genetic diversity, particularly for a conifer (Ledig and Conkle 1983; Farjon 2013; Di Santo et al. 2022), and F1 hybrids between the two populations appear to have higher fitness proxies than the parental populations, suggesting genetic rescue may be beneficial (Hamilton et al. 2017). However, increased F1 fitness may be the result of heterosis, in which excess homozygosity of deleterious recessive alleles is reduced (Williams and Savolainen 1996; Tallmon et al. 2004; van de Kerk et al. 2019). Given this, it is unclear whether increased hybrid fitness would persist in future generations of hybrids backcrossed with either parental population. If the parental populations have evolved partial reproductive isolation as a result of ecological speciation, F2s and backcrossed individuals may ultimately have lower fitness despite heterosis in F1s, limiting the value of inter-population genetic rescue (Walter et al. 2020; Christie et al. 2022).

In this paper, we investigate whether there is evidence for incipient ecological speciation between Torrey pine populations by testing whether loci with reduced admixture in F1 hybrids are associated with fitness-related phenotypic differences. We address three main questions: 1) Are there genomic signatures of reduced admixture between the two Torrey Pine populations? 2) If so, are alleles with reduced admixture linked to phenotypes or functions that indicate they may be involved in local adaptation and underlie extrinsic postzygotic isolation? 3) What are the implications for conservation of this endangered species – would genetic rescue be beneficial or should these two populations be managed as separate taxonomic groups?

## Methods

### Multigenerational common garden

Prior to 1960, two plots of 20 mainland and 20 island Torrey pine trees were established adjacent to each other at the USDA Horticultural Field Station (now the Scripps Institute) in La Jolla, CA (Haller 1967; Hamilton et al. 2017). In 2004, open-pollinated cones from 10 mainland and 16 island trees were sampled in the plots, yielding a total of 643 seeds. In 2006, seeds were planted and grown in a greenhouse at the USDA-Forest Service Pacific Southwest Station, Institute of Forest Genetics, Placerville, CA. As seedlings emerged, one cotyledon was clipped from each plant for genetic analyses and identification of seedlings’ ancestry (island, mainland, or F1 hybrid) using two population-specific allozyme markers (Ledig and Conkle 1983). Of the 643 seeds planted, 128 were genetically confirmed as being of pure mainland ancestry, 134 were identified as being of pure island ancestry, and 381 were identified as mixed ancestry (F1 hybrids) (Ledig and Conkle 1983; Hamilton et al. 2017). These F1 hybrids solely reflected crossings of island females with mainland pollen donors, indicating hybrids were the product of unidirectional gene flow between parental populations. In 2007, a subset of 360 seedlings representing each ancestry (n=120 each of island, mainland, and F1 hybrids) were planted following a randomized complete block design with six replicate blocks at the common garden site in Montecito, CA (34.42963, -119.56396). Each of the 360 trees planted were measured for four quantitative traits once a year (Appendix S3), including tree height in cm (measured between 2008 and 2018), the number of conelets or first-year cones produced (measured between 2013 and 2018), the number of immature cones or second-year cones produced (measured between 2015 and 2018), and the number of mature or third-year cones produced (measured between 2015 and 2018). These traits were selected as they provide multiple proxies of fitness, incorporating both growth and reproductive output needed to examine the impact of interpopulation gene flow to Torrey pine fitness (see below). Given that selection likely acts on each reproductive stage and there was limited correlation across traits over time (see below), we analyzed each cone production trait separately using all available years. For full details about the common garden experiment see Hamilton et al. (2017).

To understand the effects of climate on fitness, we compared climatic variables from the two native Torrey Pine populations and the common garden site, extracting data from climate rasters from *BioClim* version 2.1 for the years 1970-2000 at 30 second resolution (Fick and Hijmans 2017) using the R package *terra*, version 1.7.71 (Hijmans 2024) in R 4.3.2 (R Core Team 2023). The common garden site is a novel environment compared to either native site, and is more similar to the island site along PC1 (Appendix S2). However, because the island site generally experiences more moderate conditions (lower temperatures and more precipitation during the warmest quarter, Appendix S1 and S2), and because the severity of the summer drought is the primary stressor in Mediterranean climates and is likely to be a more important factor in adaptation (Granda et al. 2014; Nardini et al. 2014; Riordan et al. 2016; DeSilva and Dodd 2020), we considered the common garden site to be more similar to the native mainland site.

### DNA extraction and reduced representation sequencing

During the summer of 2016, needle tissue was collected from the 262 individuals surviving of the 360 seedlings originally planted within the common garden (Hamilton et al. 2017), including 89 pure mainland, 88 pure island, and 85 F1 hybrid trees. Following collection, needles were dried and maintained on silica gel until genomic DNA could be extracted using approximately 20 to 35 mg of dried needle tissue based on a modified version of the CTAB protocol (Doyle and Doyle 1987). In this modified version, rough mixing (e.g., vortex) was replaced with slow manual shaking of samples to reduce DNA shearing. The concentration and purity of extracted DNA was evaluated for each sample separately using a NanoDrop 1000 Spectrophotometer (Thermo Scientific). Overall, purity ratios averaged 1.86 and 2.21 for 260/280 and 260/230 respectively, with concentrations of DNA ranging from 153.78 to 4,463.31 ng/µL (average = 1,138.41 ng/µL). Following extraction, DNA concentration was standardized to 85 ng/µL for all 262 samples and reduced-representation sequencing libraries were prepared using the protocol described in Di Santo et al. (2022). Once constructed, all libraries were pooled and sent to the Genomic Sequencing and Analysis Facility (GSAF; Austin, TX) for single-end sequencing (1x100 bp) on 5 lanes of an Illumina HiSeq 2500. Prior to sequencing, the pooled library was size-selected for fragments within the range of 450-500 bp.

### Reference-guided assembly and calling of genetic variants

Raw sequenced libraries were demultiplexed using *ipyrad* version 0.9.12 (Eaton and Overcast 2020). To reduce the probability of assigning a read to the wrong individual, *ipyrad* was parameterized to allow only a single mismatch in the barcode sequence. Raw reads were subsequently filtered for quality using *dDocent* version 2.8.13 (Puritz et al. 2014a,b). Within *dDocent*, using *fastp* (Chen et al. 2018), base pairs with PHRED score below 20 at the beginning and end of reads and Illumina adapter sequences were removed, and reads were trimmed once the average PHRED score dropped below 15. Lastly, reads where more than 50% of base pairs had a PHRED score below 15 were discarded. Cleaned reads were mapped to the *Pinus taeda* (loblolly pine) draft genome version 2.0 (GeneBank accession: GCA_000404065.3) using the *BWA-MEM* algorithm (Li 2013) with default values, except for the mismatch penalty (-B, default: 4) and the gap open penalty (-O, default: 6), which were relaxed to 3 and 5, respectively, to account for additional genomic differences that may have evolved between Torrey pine and closely related loblolly pine. Genetic variants were called from sequence alignments using the *SAMtools* and *BCFtools* pipeline (Li 2011). Specifically, we used the multiallelic and rare-variant calling model (-m) with no prior expectation on the substitution rate (--prior 0). In total, the pipeline identified 44,493,992 genetic variants, including single-nucleotide (SNP) and insertion/deletion (INDEL) polymorphisms, that were subjected to quality filtering. First, variants with a genotype quality (GQ) < 20 and a genotype depth (DP) < 3 were marked as missing. Next, variants with PHRED scores ≤ 20 (QUAL), minor allele counts (MAC) < 3, minor allele frequencies (MAF) < 0.01, genotyping rates across individuals < 0.8, average depth across individuals > 23 (based on the equation from Li 2014), and inbreeding coefficients (F_IS_) < -0.5 or > 0.5 were filtered out of the raw data. Of the initial 262 genotyped individuals, 53 were removed from subsequent analyses as they exhibited greater than 50% missing values. In total, following the removal of INDELs and SNPs with 3 or more alleles, 11,379 biallelic SNPs across 209 individuals, including 68 pure island, 75 pure mainland, and 66 F1 hybrid individuals were identified and used for analysis.

### Parental population hybridization simulations

To evaluate whether barriers to gene flow may have evolved between Torrey pine populations, we quantified genome-wide admixture in F1 hybrids, measured as locus-specific observed heterozygosity (H_O_), specifically searching for SNPs with reduced heterozygosity. Using a customized R script (R version 4.1.3) relying on R packages *adegenet* version 2.1.5 (Jombart 2008; Jombart and Ahmed 2011) and *hierfstat* version 0.5.10 (Goudet and Jombart 2021), we simulated the distribution of H_O_ values for each of the 11,379 SNPs expected in the F1 hybrids under the null hypothesis that H_O_ at any locus would be based on the frequency of alleles in the island and mainland population. To do so, we used a 2-step approach within the function *hybridize()* that simulates hybridization between two populations. First, allele frequencies are derived from genotypes specific to each parental population. Then, hybrid genotypes are generated by sampling gametes in each of these populations using a multinomial probability distribution. SNP-specific null distributions were produced by repeating this process 1000 times, estimating locus-specific H_O_ each time from simulated hybrid genotypes using the function *basic.stats()*. To mirror the empirical SNP data set, we simulated 11,379 genotypes for 66 F1 hybrids within each iteration, using allele frequencies derived from 68 and 75 pure island and pure mainland individuals, respectively.

To identify SNPs that may exhibit reduced heterozygosity in F1s relative to expectation, we compared simulated null H_O_ distributions (expected H_O_) with H_O_ values estimated from empirical F1 hybrid genotypes (observed H_O_). For each locus, we computed a one-sided p-value, defined as the probability that an expected H_O_ value is equal or lower than the observed H_O_ value at this locus. All p-values were corrected for multiple testing using Benjamini and Hochberg’s (Benjamini and Hochberg 1995) False Discovery Rate procedure implemented within the R function *p.adjust()*. SNPs that exhibited significantly reduced heterozygosity relative to expectation based on parental allele frequencies (FDR < 0.1) were classified as potential candidate regions of the genome associated with the evolution of barriers to gene flow between island and mainland populations of Torrey pine.

### Functional plant ontology (PO) and gene ontology (GO) annotation

SNPs exhibiting reduced heterozygosity relative to expectations were functionally annotated to determine whether their functions, processes, or anatomical and temporal expressions may be important to Torrey pine fitness using *blastx* 2.9.0+ (Altschul et al. 1990, 1997; Camacho et al. 2009) and a combination of customized R scripts. First, a region of 5,000 bps (5 kbps) from *Pinus taeda* draft genome version 2.0 (GenBank accession: GCA_000404065.3) was extracted 2,500 bps before and after each target locus using *samtools* (Li et al. 2009). If a SNP was within the first 2,500 bp of a scaffold, the first 5,000 bps were extracted. If a scaffold harboring a SNP was shorter than 5 kbps, the whole scaffold length was used instead. Extracted sequences were then blasted against the TAIR10 peptide blastset (TAIR10_pep_20101214_updated). Homology between query and blasted sequences were assessed using *blastx* default parameters, an expectation value threshold of 10^-3^ (-evalue 0.001), and a maximum number of database hits of 5 (-max_target_seqs 5). Lastly, mapping of gene ontology (GO) and plant ontology (PO) terms onto annotated sequences was performed in R using a customized script and GO (ATH_GO_GOSLIM.txt.gz) as well as PO (grow.txt.gz) annotations available online. All databases mentioned throughout this section are available from https://www.arabidopsis.org/download/. While neither conifer- nor pine-specific, using the TAIR database provides a more exhaustive set of GO and PO annotations to associate with reduced admixture than is available for conifers. This database remains the most complete with respect to plant gene functions and shares orthologous and homologous sequences with pine that will aid in determining functional categories critical to plant fitness (e.g., Eckert et al. 2010). Functional gene annotation is limited for conifers, particularly for genes involved in reproduction and development (De La Torre et al. 2020). Furthermore, some genes involved in flowering in angiosperms pre-date their divergence with gymnosperms (Moyroud et al. 2017; Liu et al. 2018; De La Torre et al. 2020). However, because of the evolutionary distance between *Arabidopsis* and Torrey pine, the annotations will be best understood within broad functional categorizations.

Following the same procedure as above, we also annotated 5 kb regions around 500 randomly selected SNPs of the total 11,379 called to investigate whether the set of candidate loci for reproductive isolation was enriched or depleted for specific functions or processes using Fisher’s exact test for count data as implemented in the R function *fisher.test()*. For each annotation (GO or PO), we constructed a two by two contingency table recording the number of successes (the number of times an annotation was observed) and failures (the sum of successes across all annotations minus the number of successes for the annotation of interest) in the candidate and the random set of loci, and tested the null hypothesis of independence of rows (candidate, random) and columns (success, failure). When an annotation was absent from one of the sets of loci, we considered the number of successes associated with that annotation and set of loci to be zero. Resulting p-values were corrected for multiple testing within each annotation data set (GO or PO) and category within annotation data sets (for GO: biological process, molecular function, and cellular component) using Benjamini and Hochberg’s (Benjamini and Hochberg 1995) False Discovery Rate procedure implemented within the R function *p.adjust()*. We assumed an annotation to be either enriched or depleted within a particular set of loci when the false discovery rate associated with the odd ratio for that annotation was inferior to 10% (FDR < 0.1).

### Correlation analysis between measured phenotypes

To limit redundancy across traits and ensure each phenotype largely reflects unique inter-individual variation, we conducted a repeated measures correlation analysis in R based on temporal measurements taken on all 209 samples using the package *rmcorr* (Bakdash and Marusich 2022). Significance of estimated correlation coefficients was evaluated considering a threshold α = 0.05. A correlogram with correlations between all pairs of phenotypes was generated using the R package *corrplot* (Wei and Simko 2021) and is shown in Appendix S4. While some correlation coefficients were significant, no phenotypes were highly correlated (r ≥ 0.6). Consequently, all four temporally assessed traits – tree height in cm, the number of conelets (first-year cones), the number of immature cones (second-year cones), and the number of mature cones (third-year cones) produced – were kept for subsequent analyses.

### Genome-wide association analysis (GWAS)

To assess whether genotypic variation at select SNPs may be associated with variability in phenotypes measured across all available years data (Appendix S3), we conducted a repeated-measures genome-wide association analysis based on all 209 samples and 11,379 genetic variants using the R package *RepeatABEL* (Rönnegård et al. 2016). For each trait, the function *preFitModel()* was used to fit a linear mixed model to the data without including SNP effects to estimate variance components for the trait, including a random polygenic (based on a genetic relationship matrix, Appendix S5) and permanent environmental effect. Including these effects (computed and assessed internally within the function) accounted for genetic relatedness among individuals and repeated annual measures within the model. In addition, “block” (spatial position of a sample within the common garden) and “year” were considered as random variables within the model. Following this, we used the function *rGLS()*, fitting a generalized least square model to each genetic marker given a covariance matrix (estimated using *preFitModel*), to test for associations between SNPs (considered as fixed effects) and phenotypes while correcting for the effect of random variables. Lastly, SNP-specific p-values were corrected for multiple testing using Benjamini and Hochberg’s (Benjamini and Hochberg 1995) False Discovery Rate procedure implemented within the R function *p.adjust()*. Of the 11,379 SNPs tested for association with each of the four phenotypic traits, only those with FDR < 0.1 were considered potential candidate markers underlying genotype-phenotype association.

### Evaluating fitness consequences following interpopulation genetic mixing

One locus was identified that exhibited both lower F1 heterozygosity than expected under a neutral model and was statistically associated with one or more of the four fitness or fitness-related phenotypes (hereafter referred to as the shared locus, see Results). To suggest whether this locus could be a potential candidate associated with extrinsic reproductive isolation between Torrey pine populations, we required that (1) genotype frequencies for the locus would be biased by population, with each population favoring a different homozygous genotype, and (2) individuals with non-local or admixed genotypes would exhibit reduced fitness relative to individuals with the more local genotype. As the common garden was planted in Montecito near Santa Barbara, CA, we considered mainland trees as local and island trees as non-local. To determine whether the frequency of homozygous genotypes (i.e., 0/0 and 1/1) statistically differed between parental ancestries, we counted genotype occurrences across pure island and pure mainland individuals and conducted a Pearson’s Chi-squared test with Yates’ continuity correction as implemented in the function *chisq.test()* in R for the resulting contingency table.

A linear mixed model and expected marginal means approach was used to evaluate whether average phenotypes statistically differed among genotypes by leveraging R packages *lme4* (Bates et al. 2015) and *emmeans* (Lenth 2023). For each trait, we built a model where phenotypic variation was explained by the fixed effect of individuals’ genotypes at the shared locus, the fixed effect of among-individual genetic relationships, the random effect of blocks assigned to individuals within the common garden, the random effect of year the phenotypes were measured, and the random effect of individuals themselves (to account for the repeated nature of response variables). Genetic relationships among common garden individuals were estimated using a principal component analysis implemented within the *dudi.pca()* function from the *adegenet* R package. This analysis was performed on allele frequencies of the full genomic data set (11,379 SNPs across 209 individuals), with missing values replaced by the average allele frequencies across individuals. The first two principal components were used in all four linear mixed models to summarize genetic relationships among individuals, as these components provide good proxies for among-population (PC1) and within-population genetic structure (PC2) (Appendix S6).

The *emmeans()* function, computing expected marginal means, was used for the assessment of phenotype average differences among genotypes. Briefly, the function computes genotype-specific trait averages and standard errors corrected for all effects included within linear mixed models. The function also computes adjusted p-values estimated from t-ratios calculated between all possible pairwise genotype comparisons. Ultimately, we used this statistic to infer statistical differences in average phenotypes for all traits measured between genotypes. For this analysis, 175 of the total 209 Torrey pine trees present within the full genomic data set were used, as genotypes at the shared locus were missing for 34 individuals.

### Evaluating the distribution of genetic variation across Torrey pine ancestries and SNP sets

The distribution of genetic diversity estimated as observed heterozygosity (H_O_) was compared between pure island, pure mainland individuals, and F1 hybrids for the whole genomic dataset (N=11,379 SNPs), SNPs significantly explaining variation in fitness-related traits (N=12 SNPs), and SNPs exhibiting reduced F1 heterozygosity relative to expectations (N=185 SNPs). In addition, for each ancestry separately, we quantified the number of loci across the whole genomic dataset that were fixed. A locus was considered fixed within an ancestry when all individuals of that ancestry at that locus were homozygous for either the reference or alternate allele (0/0 or 1/1). H_O_ and numbers of fixed loci were both estimated in R. While H_O_ was computed using the *basic.stats()* function implemented within the package *hierfstat*, numbers of fixed loci were computed manually.

To determine whether differences in genetic diversity exist among ancestries within SNP datasets, we performed three Dunn’s tests using the package *dunn.test* in R. This test is a nonparametric equivalent of analyses of variance and post-hoc tests, as normality for estimated observed heterozygosities within ancestries and SNP sets, assessed either visually or using Shapiro-Wilk normality test, could not be assumed. We accounted for multiple testing by correcting p-values associated with each comparison using Benjamini and Hochberg’s (Benjamini and Hochberg 1995) False Discovery Rate procedure. Two H_O_ averages were considered significantly different when the false discovery rate dropped below 10% (FDR < 0.1).

Finally, Fisher’s exact test for count data as implemented within the function *fisher.test()* in R was used to compare the number of loci that are homozygous for either reference or alternate alleles between mainland, island, and F1 individuals. We first leveraged a three by two contingency table recording the number of successes (the number of fixed loci) and the number of failures (the total number of loci minus the number of fixed loci) across all three ancestries, testing the null hypothesis of the independence of rows (island, mainland, F1 hybrid) and columns (success, failure). In addition, we used three two by two contingency tables of the number of successes and failures (defined as above) to test the null hypothesis of the independence of rows (F1 hybrid, island; F1 hybrid, mainland; island, mainland) and columns (success, failure). The p-value associated with each pairwise comparison was corrected for multiple testing using Benjamini and Hochberg’s (Benjamini and Hochberg 1995) False Discovery Rate procedure implemented within the R function *p.adjust()*. We considered a difference to be significant when the false discovery rate dropped below 10% (FDR < 0.1).

## Results

### SNPs exhibiting lower-than-expected F1 heterozygosity and their functional relevance

Comparison of simulated F1 hybrids between island and mainland individuals for 11,379 SNPs using empirical allele frequencies in each parental population revealed some loci that exhibited a higher degree of homozygosity in the F1s relative to the expectation. A comparison between observed and simulated H_O_ estimates at each locus identified 185 SNPs (FDR < 0.1) with reduced F1 heterozygosity. Of the 184 5kb-long sequences containing these SNPs (two SNPs were located on the same segment), 75 (41%) could be annotated with the TAIR 10 peptide blastset, with hits in 59 described *A. thaliana* genes. Plant ontology terms could be retrieved for 37 (63%) of the 59 gene hits, while gene ontology terms were retrieved for all 59 gene hits.

Regions of the genome that exhibited reduced F1 heterozygosity relative to expectation were associated with a variety of plant developmental stages (Appendix S7). This includes embryo development stages (e.g., C globular stage, D bilateral stage, E expanded cotyledon stage, or F mature embryo stage), vegetative development stages (e.g., 2-, 4-, 6-, 8-, 10-, 12-leaf stages, or seedling development stage), and reproductive development stages (e.g., 4 anthesis stage, petal differentiation and expansion stage, M germinated pollen stage, or L mature pollen stage). Functionally, the majority of these loci are located or active in the nucleus with molecular functions including protein binding, catalytic activity, RNA binding, as well as kinase and transferase activities (Appendix S8A, B). Biological processes associated with these molecular functions covered responses to various stimuli and stresses, and plant development (Appendix S8C). Enrichment analyses demonstrated that while no PO terms were preferentially associated with SNPs exhibiting lower heterozygosity than expected in the F1, some GO terms were significantly enriched or depleted in the latter SNP set (Figure 1, Appendix S9). Interestingly, assessment of enriched GO terms indicated the products of genes associated with these loci may be preferentially located in the plasma membrane and involved in important developmental and reproductive processes, including cell growth, plant growth, and pollination, with molecular functions associated with signaling receptor activity.

**Figure 1.**
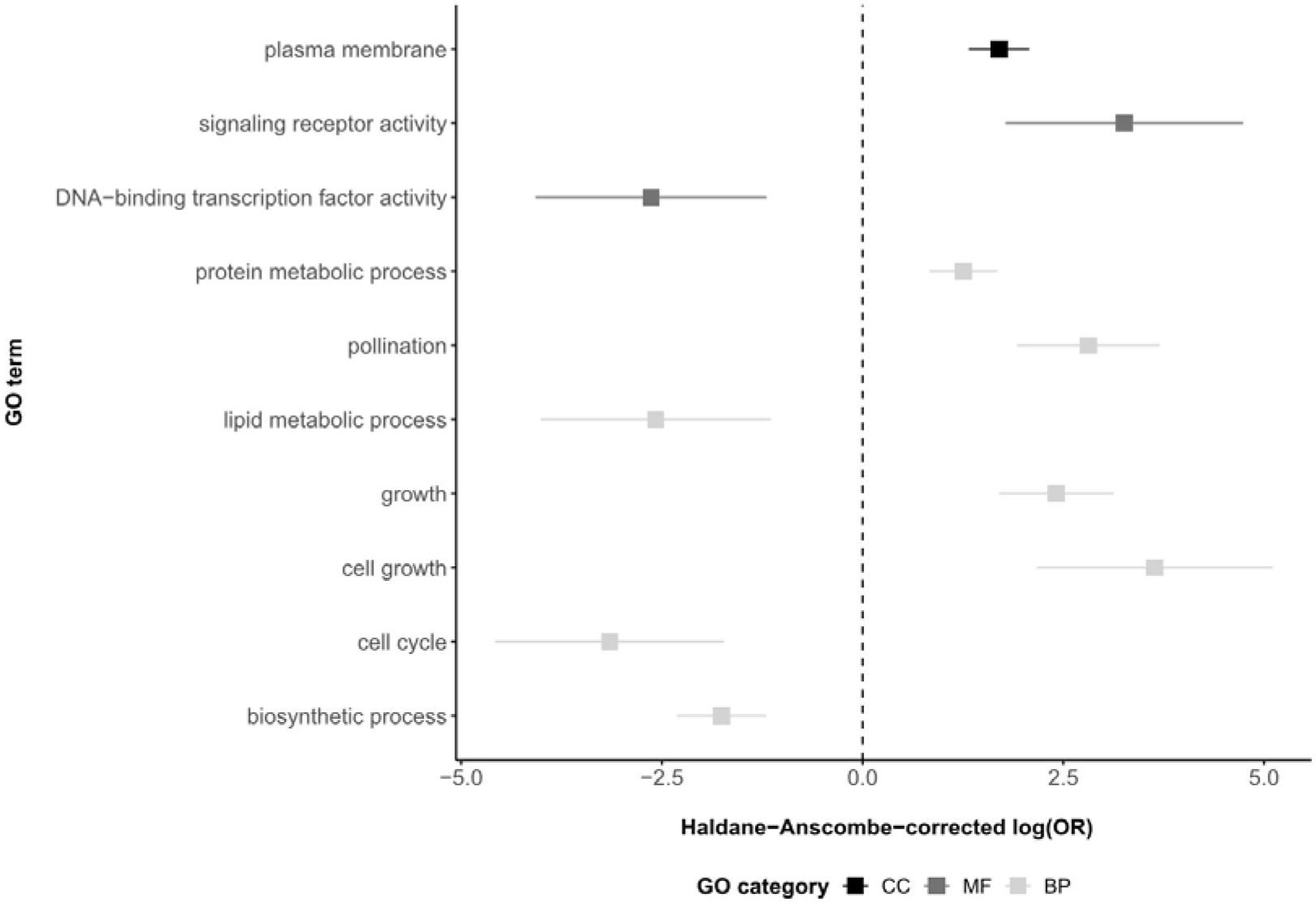
Haldane-Anscombe-corrected log-transformed odd ratios ± standard error (x axis) for significantly (FDR < 0.1) enriched (positive values) and depleted (negative values) GO terms (y axis) for the set of alleles with reduced F1 heterozygosity, including cellular component (CC, black), molecular function (MF, dark gray), and biological process (BP, light gray) terms. Here, we present Haldane-Anscombe-corrected estimates of log-transform odd ratios as the presence of zero success probabilities within the data would otherwise lead to nonsensical estimates of odd ratios (see Appendix S9 for details).

### SNPs exhibiting statistical association with fitness traits and their proxies

Genome-wide association analysis indicated alleles at several loci may play an important role in contributing to fitness variation in Torrey pine. In total, 12 SNPs were significantly associated (FDR < 0.1) with variation in phenotypes (Table 1). While 8 (67%) of the 12 SNPs were associated with a single trait, the remaining 4 were associated with two or more traits. Phenotypic values of these traits all increased with the number of derived alleles at associated SNPs except for one, locus_8778 [APFE031129984.1:16200] (Table 1), where it decreased. Interestingly, comparing SNPs associated with fitness and fitness-related traits with SNPs exhibiting low-level F1 heterozygosity identified one common locus (locus_4218 [APFE030529380.1:392010]). This locus, in addition to exhibiting a reduced degree of F1 heterozygosity (Appendix S10), was also the only locus that significantly explained phenotypic variation across all four traits measured (Table 1). Unfortunately, functional annotations for the 5-kbps region surrounding this locus were limited. While this region may be located in the mitochondrion (GO:0005739), nothing is known about its molecular function or the biological process it might be involved in.

**Table 1.** SNPs significantly associated with one or more phenotypes (FDR &lt; 0.1), including tree height (cm), number of conelets, number of immature cones, and number of mature cones. Listed are generic SNP IDs given for this study, SNP IDs retrieved from *Pinus taeda* draft genome (scaffold ID:position on the scaffold), the trait(s) each SNP is associated with, the effect of increasing the number of derived alleles on the associated phenotype(s), and the FDR of the genotype-phenotype association.

| Generic<br>SNP ID | SNP ID within<br>*P. taeda* genome | Phenotypic trait<br>association | Phenotypic<br>effect | FDR |
| --- | --- | --- | --- | --- |
| locus_4218 | APFE030529380.1:392010 | Tree height (cm) | + | 0.047 |
|  |  | Nb conelets | + | <0.001 |
|  |  | Nb immature cones | + | 0.085 |
|  |  | Nb mature cones | + | 0.037 |
| locus_12498 | APFE031619559.1:67487 | Tree height (cm) | + | 0.047 |
| locus_433 | APFE030047412.1:104020 | Nb conelets | + | 0.078 |
|  |  | Nb mature cones | + | 0.057 |
| locus_3004 | APFE030371338.1:302521 | Nb conelets | + | 0.078 |
| locus_3676 | APFE030458498.1:93076 | Nb conelets | + | 0.024 |
| locus_9630 | APFE031249435.1:61968 | Nb conelets | + | 0.011 |
|  |  | Nb immature cones | + | 0.085 |
|  |  | Nb mature cones | + | 0.027 |
| locus_13093 | APFE031688147.1:172170 | Nb conelets | + | 0.072 |
|  |  | Nb mature cones | + | 0.027 |
| locus_1164 | APFE030138697.1:151998 | Nb mature cones | + | 0.032 |
| locus_6624 | APFE030828725.1:2769 | Nb mature cones | + | 0.027 |
| locus_8019 | APFE031020621.1:50406 | Nb mature cones | + | 0.053 |
| locus_8778 | APFE031129984.1:16200 | Nb mature cones | - | 0.057 |
| locus_10668 | APFE031380592.1:30882 | Nb mature cones | + | 0.053 |

### Fitness consequences of hybridization at a locus exhibiting reduced admixture and significant genotype-phenotype association

The chi-squared test performed on homozygous genotype counts for locus_4218 [APFE030529380.1:392010] demonstrated unequal distribution of homozygous genotypes between pure parental ancestries (χ^2^_1_ = 104.1, P < 0.001, Appendix S10). Homozygotes for the reference allele (0/0) were dominant among pure island individuals (94% of all genotypes) while homozygotes for the derived allele (1/1) formed the core of pure mainland individuals (97% of all genotypes). Given this result, we considered the reference allele (0) as the island allele, and the derived allele (1) as the mainland allele.

The evaluation of expected marginal means (phenotype averages corrected for within- and among-ancestry genetic structure, blocks within the common garden, year of trait measurements, and sample ID) revealed unequal fitness across reference, derived, and heterozygous genotypes. For three out of the four phenotypic traits, individuals homozygous for the island allele exhibited reduced fitness, while individuals homozygous for the mainland allele exhibited greater fitness, with heterozygous genotypes intermediate between the two (Figure 2; Appendix S11). These traits included tree height, the number of conelets produced, and the number of mature cones produced. For the number of immature cones produced, the distribution of fitness among genotypes was slightly different. Individuals homozygous for the island allele still exhibited the lowest fitness, but fitness of individuals homozygous for the mainland allele and heterozygous individuals were the highest, with no significant differences between groups. We considered the mainland allele (1) as the local, and the island allele (0) as non-local because both the common garden site and the mainland population experience warmer and drier summers (Appendix S1). Our results thus indicate that carrying at least one copy of the local allele increased tree fitness, with a direct additive relationship based on the dosage of mainland allele to fitness for tree height (cm), the number of conelets produced, and the number of mature cones produced (Figure 2).

**Figure 2.**
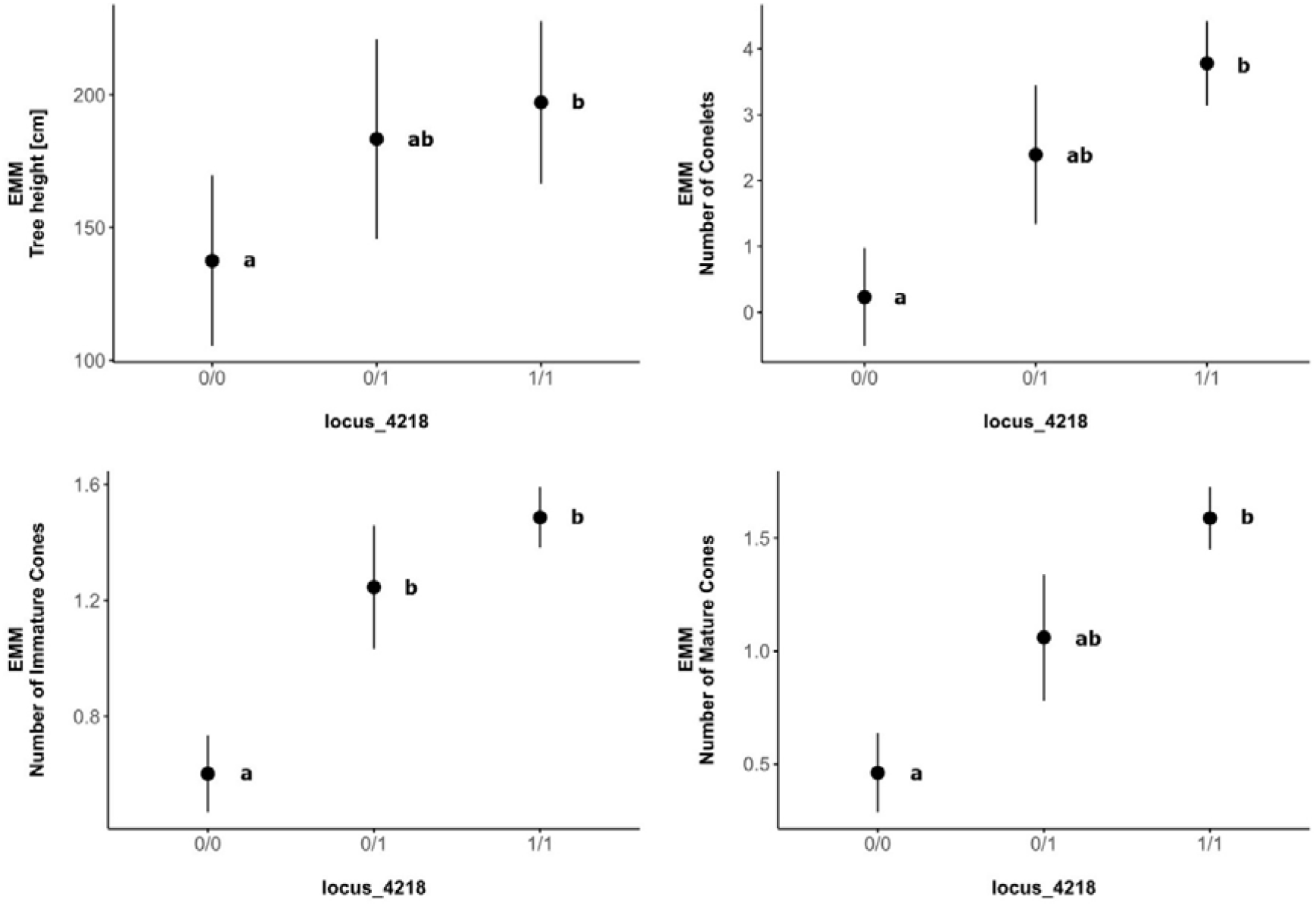
Expected Marginal Means (EMM) ± standard error of all four phenotypes investigated (y axis) given the genotype (x axis) of island, mainland, and hybrid individuals at locus_4218. Phenotypes measured include tree height (cm), number of conelets, number of immature cones, and number of mature cones. Sample sizes are 47, 5, and 123 individuals with genotype 0/0, 0/1, and 1/1, respectively. Distinct bolded lowercase letters indicate significant differences in phenotypes (adjusted P < 0.05) among genotypes (see Appendix S11 for details). Based on genotype frequencies (see Results), 0 was defined as the island allele, and 1 as the mainland allele.

### Distribution of genetic variation across island, mainland and F1 hybrid individuals

For the whole genomic dataset (N=11,379 SNPs), estimates of observed heterozygosity were similar across ancestries (H_O_ = 0.229, 0.230, and 0.230 for F1 hybrid, island, and mainland individuals, respectively; Figure 3). However, despite almost-identical average observed heterozygosities, the number of loci homozygous for either the reference or the derived allele (1/1 or 0/0) varied significantly among ancestries (Fisher’s exact test, P < 0.001), with island and mainland individuals exhibiting four times greater number of fixed alleles than that estimated for F1 hybrids (Appendix S12). While these results may seem contradictory, they can be explained by a proportionally higher number of loci with extremely low heterozygosity in the F1s relative to island or mainland individuals (Appendix S13).

**Figure 3.**
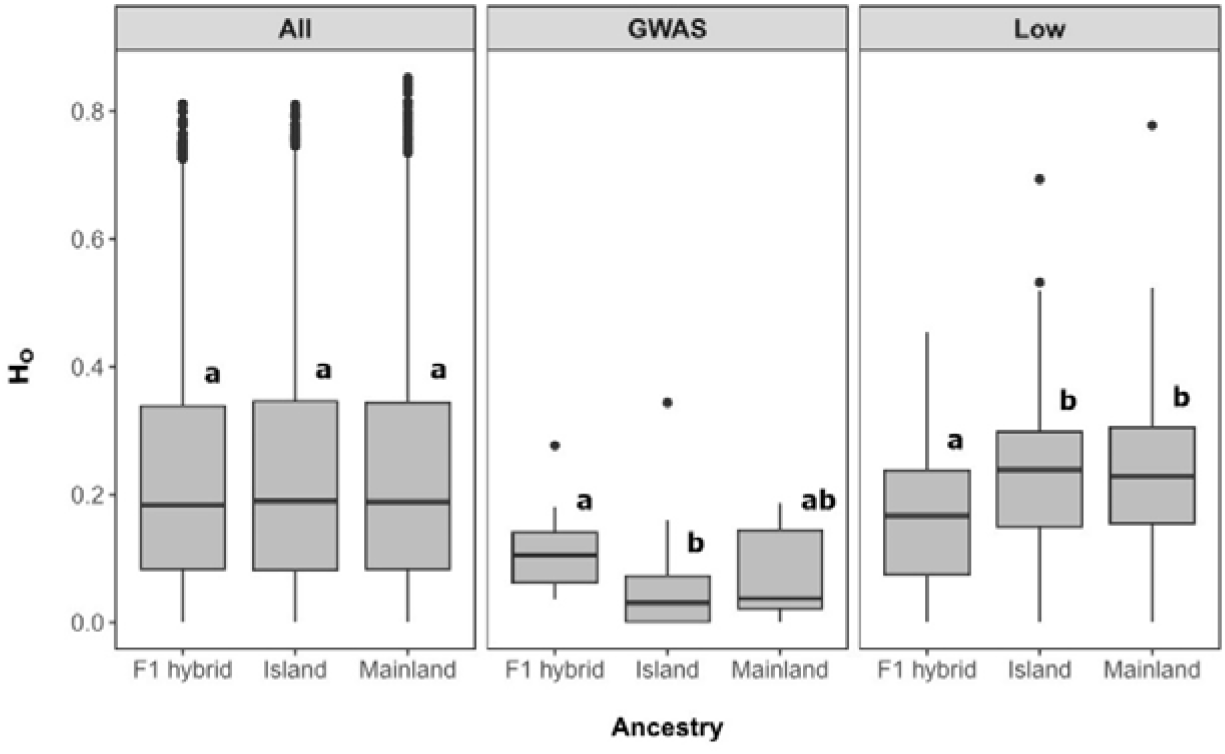
Distribution of observed heterozygosity estimates across ancestries (F1 hybrid, Island, and Mainland) for three SNP sets (All, GWAS, and Low). Distinct lowercase letters indicate significant difference in observed heterozygosities (FDR < 0.1) among ancestries (see Appendix S12 for details). All: data set containing all retained SNPs after filtering (11,379 SNPs across 209 individuals), GWAS: data set containing SNPs exhibiting a significant genotype-phenotype association (12 SNPs across 209 individuals), Low; data sets containing SNPs exhibiting reduced heterozygosity relative to expectations (185 SNPs across 209 individuals).

Average observed heterozygosity estimated for loci exhibiting reduced F1 heterozygosity for hybrids (0.159) was significantly lower than that for island (0.230) and mainland (0.232) individuals (Figure 3). If they were to contribute to extrinsic reproductive isolation, we would expect heterozygosity at these loci to be much lower in parental populations than in F1s as well as to be highly differentiated between island and mainland populations due to divergent selection promoting fixation of distinct alleles. Because average heterozygosity in F1 hybrids at these loci was lower than that found for both island and mainland individuals and given that these loci did not only comprise those highly differentiated between parental populations (Appendix S14), our results suggest intrinsic genetic incompatibilities may potentially have evolved between populations, with some allelic combinations following hybridization leading to reduced hybrid fitness.

Finally, for SNPs associated with fitness proxies, we observed exceedingly low genetic variability (Figure 3), with average estimates of heterozygosity for F1, island, and mainland individuals evaluated at 0.113, 0.066, and 0.071, respectively. These results indicate that all three groups have reduced genetic variation available for selection to act upon. However, F1 hybrids did exhibit significantly greater heterozygosity for genetic variants associated with fitness when compared to island individuals.

## Discussion

Local adaptation is common in tree species (Savolainen et al. 2007; Lind et al. 2017, 2018; Gugger et al. 2021), but it is unclear when and how population differentiation may drive the evolution of reproductive isolation. Understanding whether reproductive isolation has developed between two populations is essential to species management, particularly where intra-specific gene flow may be a beneficial source of genetic variation or may be a hindrance, leading to outbreeding depression. In Torrey pine, previous work has shown that the island and mainland populations are genetically and phenotypically different despite some recurrent gene flow, consistent with adaptation to differing environments (Hamilton et al. 2017; Di Santo et al. 2022). Furthermore, asymmetric gene flow between populations when grown together provides some evidence for the evolution of reproductive isolation (Hamilton et al. 2017). Together, this suggests that the two Torrey pine populations could be at an early stage of speciation. Here, we tested for further evidence for the evolution of reproductive isolation by identifying loci exhibiting less heterozygosity in F1 hybrids than expected given allele frequencies in parental populations, suggesting selection against heterozygotes. One locus that was highly diverged among populations and showed signatures of selection against F1 hybrids was associated with fitness differences in the common garden, and could be a candidate locus for ecologically-driven reproductive isolation between island and mainland Torrey pine populations. Overall, loci with low heterozygosity in F1 hybrids were enriched for functions important in reproduction, such as growth and pollination, suggesting these could be the mechanisms underlying the evolution of reproductive isolation between the two populations.

While demographic modeling has provided evidence for historic speciation with gene flow in pines (Menon et al. 2018; Bolte et al. 2022), the specific molecular mechanisms underlying reproductive isolation in conifers are not well understood (Bolte and Eckert 2020). We identified loci that may be involved in the evolution of reproductive isolation between two populations, including pollination- and growth-related genes, and one locus associated with fitness differences expected when hypothesizing local adaptation. In combination with asymmetric gene flow, this suggests that both intrinsic and extrinsic barriers could be contributing to reproductive isolation in Torrey pine. These results thus identify potential molecular mechanisms underlying the process of reproductive isolation in a critically endangered pine and suggest that differential adaptation could be a driving force, which is critical information required for optimal conservation management of the species.

### One locus shows patterns suggestive of extrinsic postzygotic reproductive isolation

We identified a locus that is divergent between the two populations and is also associated with fitness differences (locus_4218 [APFE030529380.1:392010]). Individuals with the derived “mainland” variant of this locus had higher growth and reproduction in the mainland common garden than individuals homozygous for the reference “island” allele, consistent with local adaptation to the hotter, drier environments shared by the mainland population and the common garden environment (Appendix S1, S2). Torrey Pine is likely a relictual endemic species that once had a larger range and became restricted to coastal climates as climate became warmer and drier (Axelrod 1967; Williams et al. 2008), and thus the island population may have a climate more similar to its ancestral climate. Demographic modeling suggests that the two modern-day Torrey Pine populations formed after an ancestral population split, forming a slightly larger island population (N_e_ = 2305) and smaller mainland population (N_e_ = 1715) approximately 1.2 million years ago, with gene flow occurring following divergence (Di Santo et al. 2022). It is possible that the mainland population has undergone additional evolutionary change following divergence to adapt to the combined stresses of warm and drier summers associated with the mainland environment, and that this locus is a candidate for local adaptation and divergence of the mainland population. The differentiation between populations for this allele could result from differential selection if heterozygous hybrids have lower fitness in both natural populations relative to individuals that are homozygous for the local variant. If this locus experiences differential selection between the two Torrey pine populations, it could contribute to reproductive isolation. While our results suggest that this locus may be beneficial in the hotter, drier summers experienced by mainland environments relative to the island population (Appendix S1), a reciprocal transplant of the two populations and their F1 hybrids would be required to determine whether the non-local variant decreases fitness in both natural populations.

Reduced average heterozygosity at this locus in approximately 10-year-old F1 hybrids (0.028 compared to a neutral expectation of 0.936) suggests selection against heterozygotes with one copy of the island and one copy of the mainland allele, or that this locus is linked to such a gene under selection. Reduced heterozygosity could result from intrinsic prezygotic or postzygotic barriers (e.g., incompatibilities affecting pollination, germination, or development of seedlings), extrinsic environmental selection in the common garden in the following ten years, or both. In the common garden, heterozygotes were intermediate in fitness metrics between island homozygotes with lower fitness and mainland homozygotes with higher fitness (although only the number of immature cones was significantly different between island homozygotes and heterozygotes, Figure 2). Our evidence suggests that selection against heterozygotes occurred at early developmental stages, reducing the expected number of heterozygous individuals, and also that the surviving heterozygous individuals in the common garden had intermediate fitness across multiple years of growth and reproduction measurements. This may seem contradictory, but it could indicate that this allele has different fitness effects at different developmental stages, or that pleiotropic interactions with other alleles result in selection against heterozygotes only at certain developmental stages or in combination with specific variants of other alleles.

While there is limited functional annotation for genomic regions proximal to this locus, it does have the annotation “mitochondrion” (GO:0005739). As heterozygotes are present, it is probably not encoded in the mitochondrion and may represent ancient paralogy between nuclear and mitochondrial genomes. Alternatively, the locus could also be part of a nuclear mitochondrial gene present in the nucleus following mito-nuclear gene transfer, with its product interacting with mitochondria. Consequently, the locus could be involved in cytoplasmic incompatibility between the maternally-inherited mitochondrion and nuclear paternal genes (such as those involved in pollen tube growth) (Turelli and Moyle 2007; Rieseberg and Blackman 2010), or with the chloroplast, which is paternally inherited in pines (Neale and Sederoff 1989; Mogensen 1996).

Local adaptation is generally polygenic in forest trees (Yeaman et al. 2016; MacLachlan et al. 2021), suggesting ecological speciation may also have a broad genetic basis (Rose et al. 2018; Schluter and Rieseberg 2022). However, we only find one locus showing evidence consistent with ecological speciation. Because we used ddRADseq data and only sampled a fraction of the genome of Torrey pine, it is likely that other loci underlying reproductive isolation and local adaptation exist, but were not captured in this study. Nonetheless, the fact that we identified a locus linked to both reproductive isolation and fitness despite this limitation suggests that the pattern may be more prevalent throughout the genome and could be better assessed using whole-genome sequence data.

### Functions of genes exhibiting reduced heterozygosity

If loci exhibiting reduced heterozygosity in F1s are “speciation genes” underlying hybrid incompatibilities, differential adaptation, or both (or linked to involved genes), they should be enriched for related functions (Rieseberg and Blackman 2010; Wright et al. 2013; Walter et al. 2020; Schluter and Rieseberg 2022). Intrinsic barriers are likely to be related to pollination or reproduction, and extrinsic barriers may be involved in traits under divergent selection, such as abiotic or biotic conditions. We found that highly differentiated loci were located near genes that were enriched for a wide range of functions (Figure 1, Appendix S9). Similar to previous studies, enriched functions included those related to pollination and development (“growth”, “pollination”, “cell growth”) (Rieseberg and Blackman 2010; Leroy et al. 2020). These regions may be involved in intrinsic reproductive barriers, such as pollination incompatibilities associated with female cones or failed embryo development post-fertilization (McWilliam 1959; Fernando et al. 2005). Other enriched functions include “plasma membrane”, “signaling receptor activity”, or “protein metabolic process”. Given these broad categorizations it is difficult to determine whether these genes could be involved in adaptation to contrasting environments, particularly as only one differentiated gene was associated with fitness differences in the common garden. Previously, island seedlings were found to germinate later and have a reduced growth rate relative to mainland seedlings (Hamilton et al. 2017). The enrichment of loci with the “growth” and “cell growth” GO terms may be related to the evolution of differential growth rates between parental populations.

However, it is also possible that the enriched functions are associated with fitness in traits or environmental conditions that were not measured in the common garden. Because the loci with reduced heterozygosity in F1 hybrids were investigated after ten years of growth in the garden, we cannot determine the stage at which selection against heterozygotes occurred in order to distinguish between intrinsic and extrinsic postzygotic barriers. Crossing experiments and monitoring of hybrids produced between the two populations would provide a more direct approach to determine the developmental stage at which potential barriers act (Christie et al. 2022).

### Conservation implications

Similar to previous studies, genome-wide estimates of heterozygosity for the island and mainland populations were low (Di Santo et al. 2022), suggesting a lack of evolutionary potential. Additionally, we found in the present study exceedingly low diversity at loci putatively underlying trait variation in the species, further supporting these results.

F1 hybrids between mainland and island populations exhibit heterosis, with a higher growth rate and fecundity than either parental population (Hamilton et al. 2017). The four-fold reduction in the number of fixed alleles when compared to pure island and pure mainland Torrey pine trees, and the almost two-fold increase in average heterozygosity at loci exhibiting significant genotype-phenotype associations when compared to pure island Torrey pine trees indicate that heterosis in F1 hybrids may result from the masking of deleterious mutations. Alone, these results would suggest that natural populations could benefit from genetic rescue, where hybridization between the two populations could introduce genetic variation to alleviate genetic load. However, our results here warrant caution before introducing non-local or hybrid trees into the wild. If the natural populations have already developed reproductive isolation or are locally adapted, hybridization may result in decreased fitness that only presents in F2s or later generation hybrids (Lowry et al. 2008a; Walter et al. 2020; Christie et al. 2022), or only in field conditions where they are exposed to stresses not present in the common garden (Melo et al. 2014). Our data suggests that the island and mainland populations should be managed separately at present, but that future work should use crossing experiments and reciprocal transplants to determine whether genetic rescue could have long-term benefits to this endangered tree species.

## Conclusions

Taken together, we find evidence of asymmetric barriers to gene flow, reduced heterozygosity in genomic regions related to varying reproductive and developmental functions, and one low-heterozygosity and highly differentiated locus that associates with reduced fitness in individuals carrying one or two non-local (island) alleles for all four measured fitness metrics in Torrey Pine. These results suggest that Torrey Pine populations may be beginning to evolve reproductive isolation, and this may partly be driven by adaptation to contrasting island and mainland environments, in which case introducing non-local individuals or hybrids to the natural populations for genetic rescue may actually be harmful in the long term. In the future, whole-genome sequencing combined with reciprocal transplant experiments and the development of experimental crosses including F2s and backcrosses will be poised to identify the genes that may be under divergent environmental selection in the two populations, and/or contributing to the evolution of reproductive isolation (Schluter and Rieseberg 2022). Furthermore, understanding the extent of reproductive isolation among these two populations will inform management strategies for this rare species that balance the benefits of genetic diversity and local adaptation.

## Data accessibility

Genomic and phenotypic data generated for this study, as well as scripts used for analysis are available from Dryad: doi:10.5061/dryad.pc866t1xg.

## Author’s contributions

***L.N.D.S.*** designed the study, contributed to data collection, generated, processed, and analyzed genomic data, interpreted the data, and led the writing of the manuscript. ***A.M.*** contributed to data collection, analyzed the climate data, interpreted the results, and led the writing of the manuscript. ***J.W.W.*** contributed to data collection and edited the manuscript.

***J.A.H.*** designed the study, contributed to data collection, interpreted the results, and wrote the manuscript. ***L.N.D.S.*** and ***A.M.*** contributed equally to the study.

## Acknowledgements

Tom Ledig, USDA-Forest Service, PSW, had the foresight to establish the common garden study at the end of his career. We thank him for sharing this project with us. For their help collecting needle tissue and measuring phenotypes within the common garden, the authors would like to thank Alexis Pearson, Tyler Stadel, Courtney Canning, and Madison Snider. The authors also would like to thank John Gabbert for allowing the scientific crew to access the common garden study. Finally, the authors would like to acknowledge the Michumash and the Kumeyaay people as the traditional caretakers of the Torrey pine ecosystems. Any use of product names is for informational purposes only and does not imply endorsement by the US Government. The findings and conclusions in this publication are those of the authors and should not be construed to represent any official USDA or U.S. Government determination or policy. This work was supported in part by the U.S. Department of Agriculture, Forest Service.

