## Supplementary Material for "Genetic basis of reproductive isolation in Torrey pine (*Pinus torreyana* Parry): insights from hybridization and adaptation"

### Appendix S1. Climate values from BioClim data (1970-2000) for each of the three sites: the common garden in Montecito (“Garden”), Torrey Pines State Reserve in San Diego County (“Mainland”), and Santa Rosa Island (“Island”).

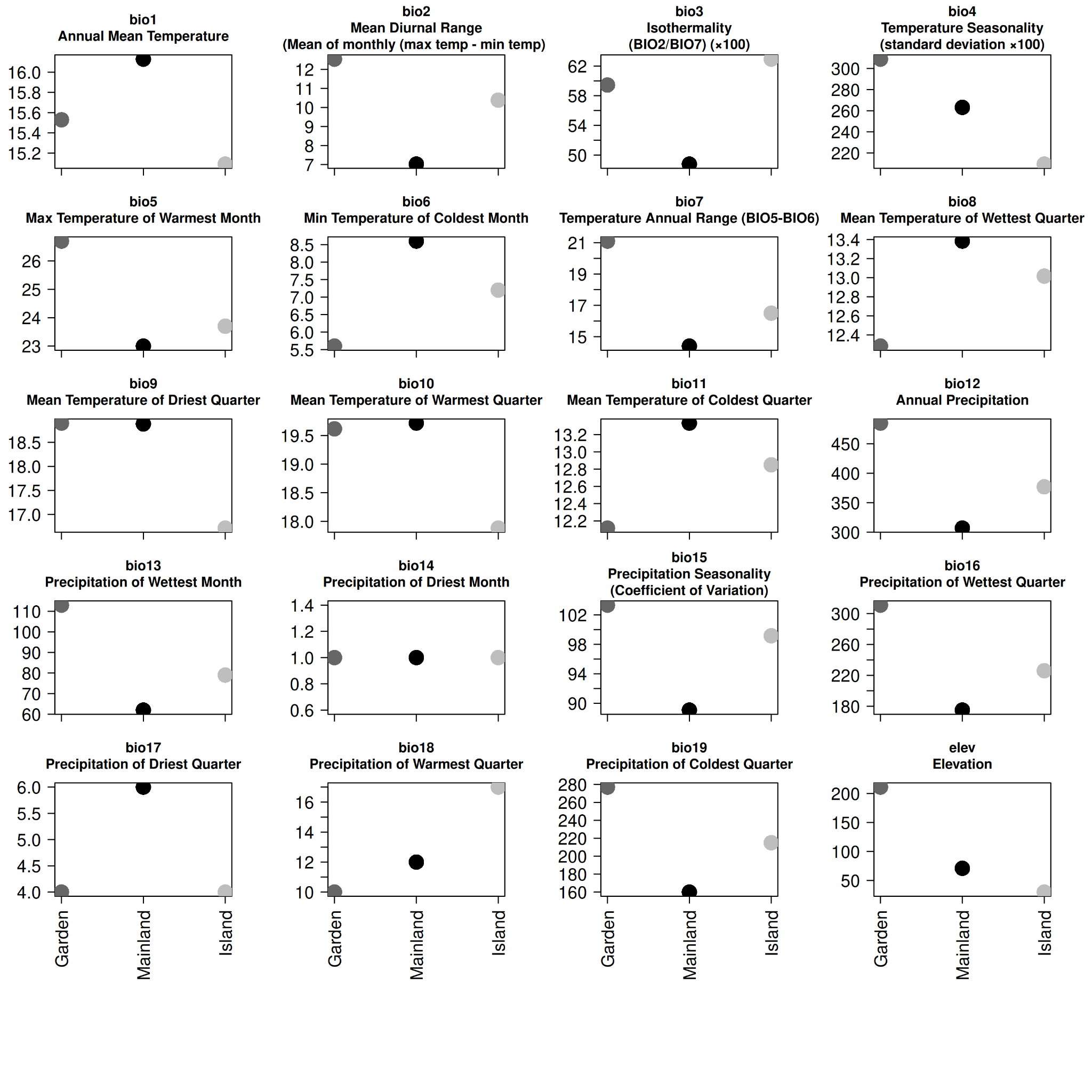

### Appendix S2. PCA of all climate values (as in S1) for each of the three sites.

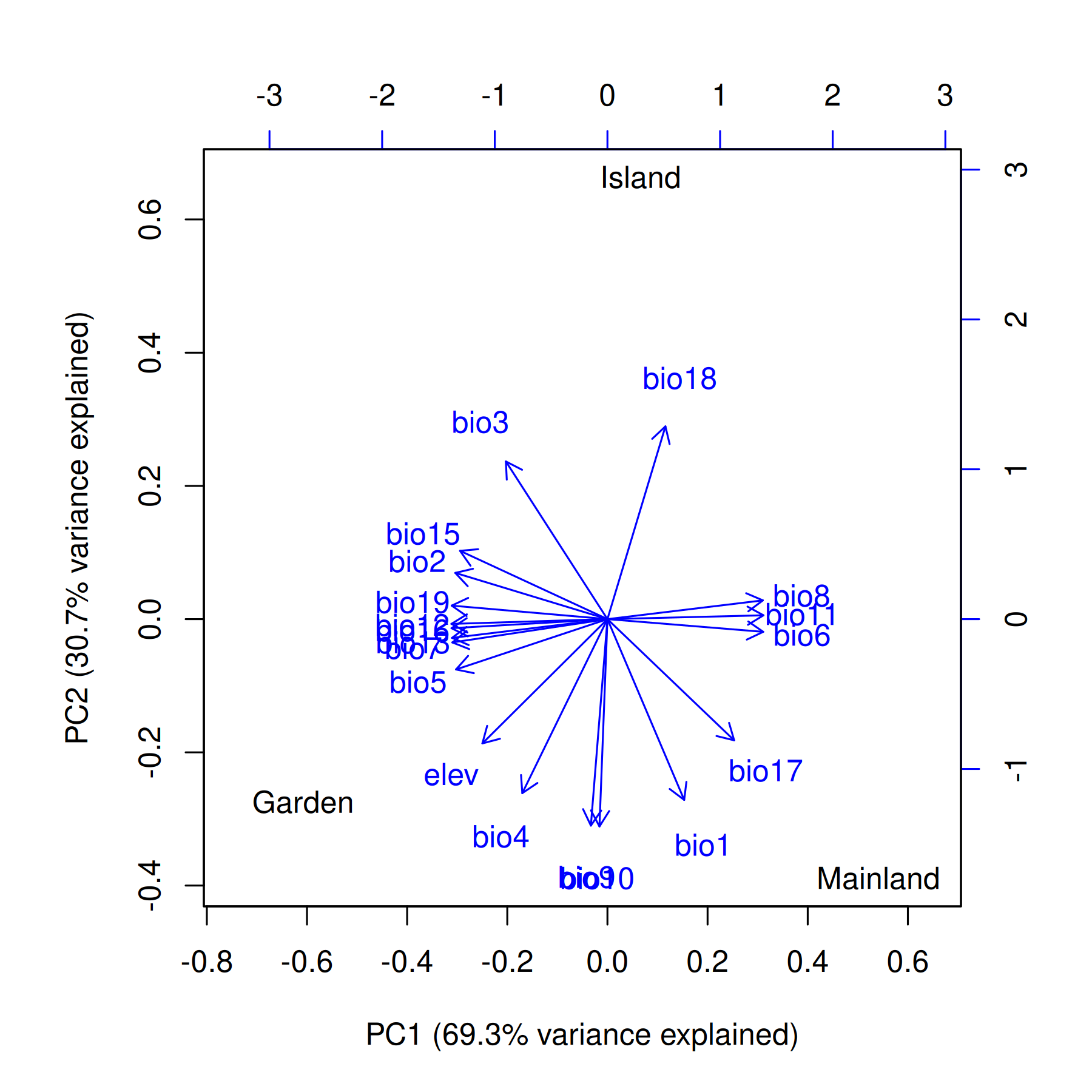

### Appendix S3. Distribution of all four phenotypes considered (y axis) for each year measurements were taken (x axis) separated by ancestries (F1 hybrid [66 individuals], island [68 individuals], mainland [75 individuals]). Phenotypes include tree height (cm), number of conelets, number of immature cones, and number of mature cones.

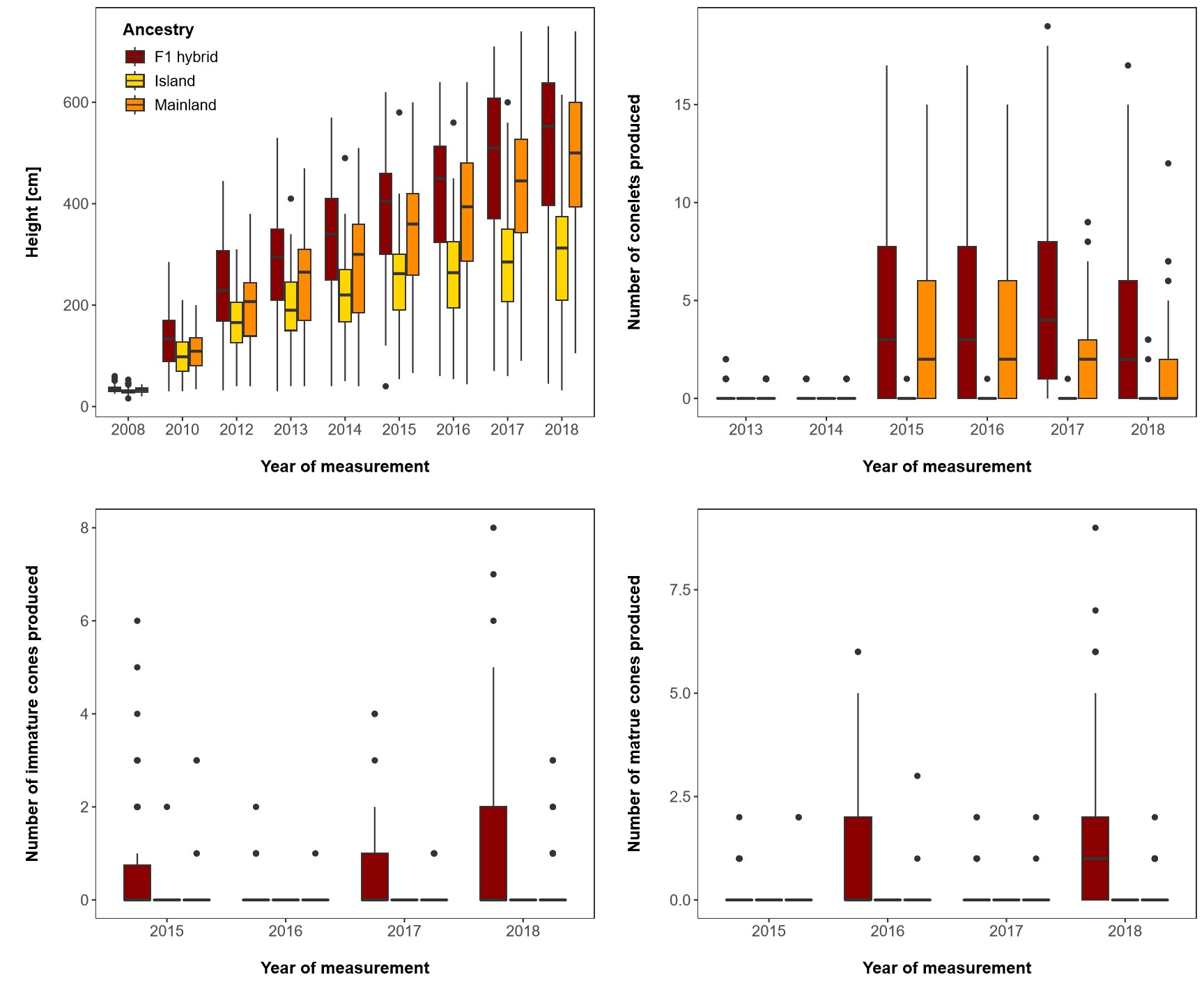

### Appendix S4. Correlogram showing repeated measures correlation coefficients among phenotypic traits, including tree height (cm), number of conelets, number of immature cones, and number of mature cones produced. The color scale represents the intensity of the correlation between two given phenotypes (the exact value is provided within each cell). Black crosses indicate non-significant correlations at α = 0.05.

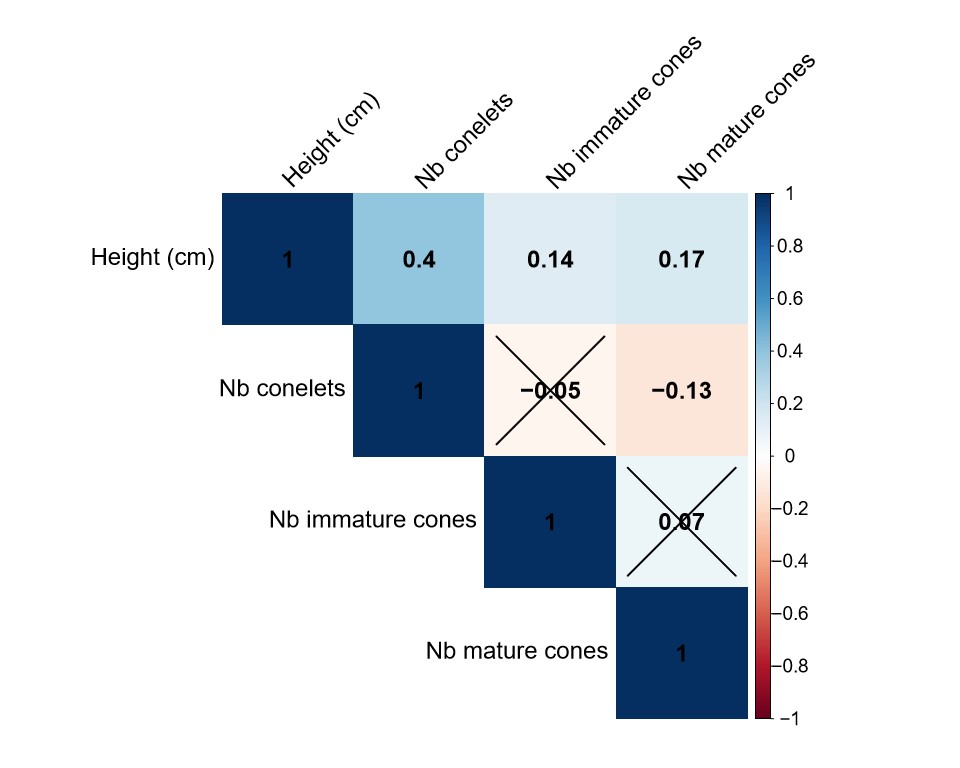

### Appendix S5. Heatmap of the genetic relationship matrix (GRM) used for genome-wide association analysis among 209 Torrey pine individuals at 11,379 SNPs. Color intensity on the white-to-red gradient scale associated with each tile represents the degree of relatedness (relatedness coefficient) between two individuals. The three colors appended to the top and to the left of the heatmap indicate the ancestry of each individual (yellow: Island [68 individuals], orange: Mainland [75 individuals], brown: F1 hybrid [66 individuals]).

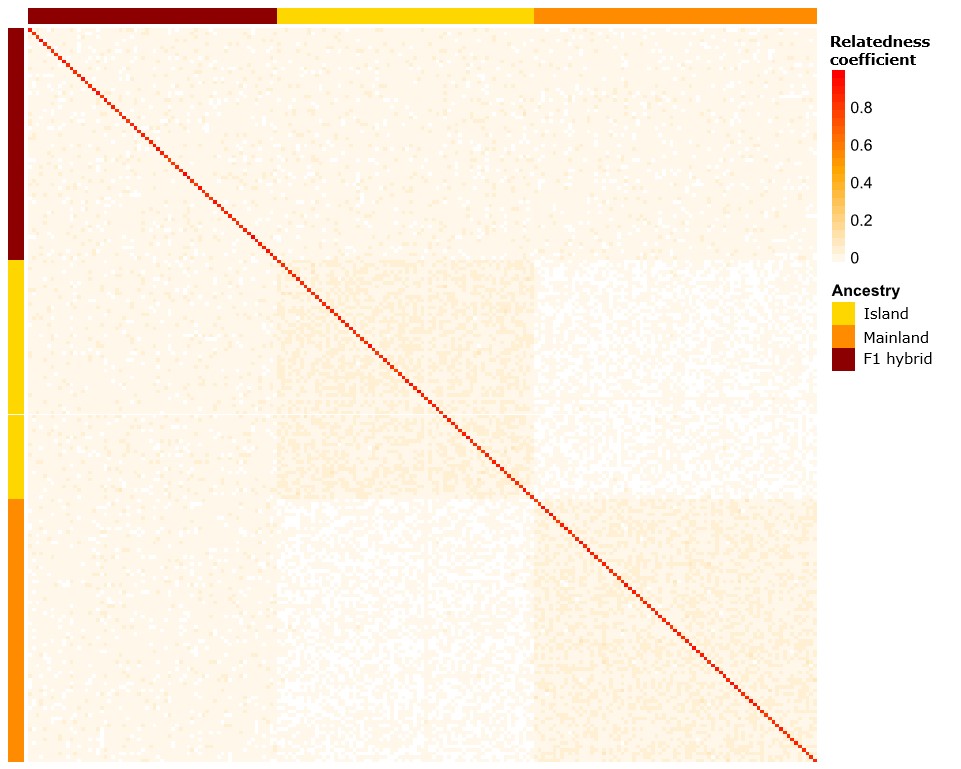

### Appendix S6. Principal component analysis using 11,379 SNPs for 209 Torrey pine samples, including 66 F1 hybrid individuals (brown), 68 island individuals (yellow), and 75 mainland individuals (orange). Variation explained by the first two principal components is given in parentheses.

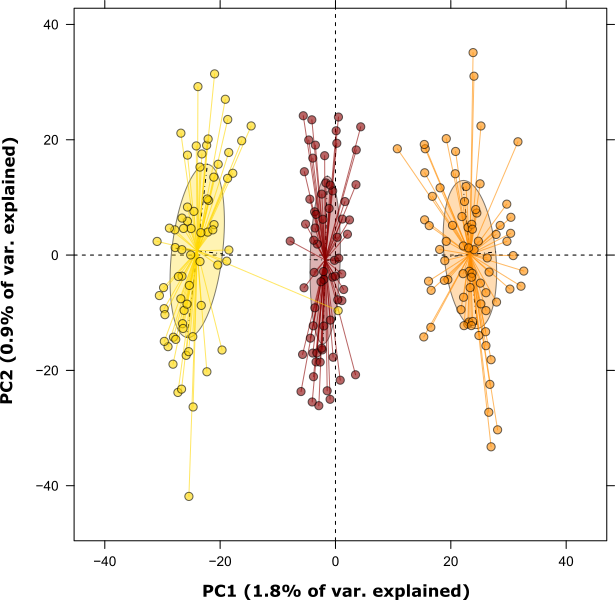

### Appendix S7. Proportion (x axis) of each plant ontology (PO) term (y axis) associated with all 75 annotated (out of 184) RI candidate segments (5kb-long sequence flanking candidate SNPs).

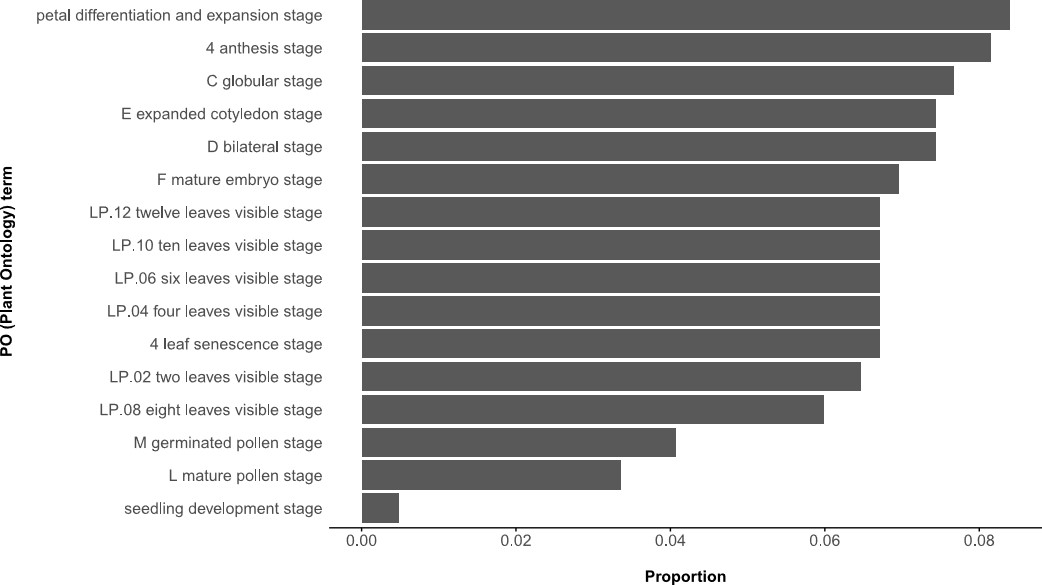

### Appendix S8. Proportion (x axis) of each gene ontology (GO) term (y axis) associated with all 75 annotated (out of 184) RI candidate segments (5kb-long sequence flanking candidate SNPs). (A) Cellular Component (CC) terms. (B) Molecular Function (MF) terms. (C) Biological Process (BP) terms.

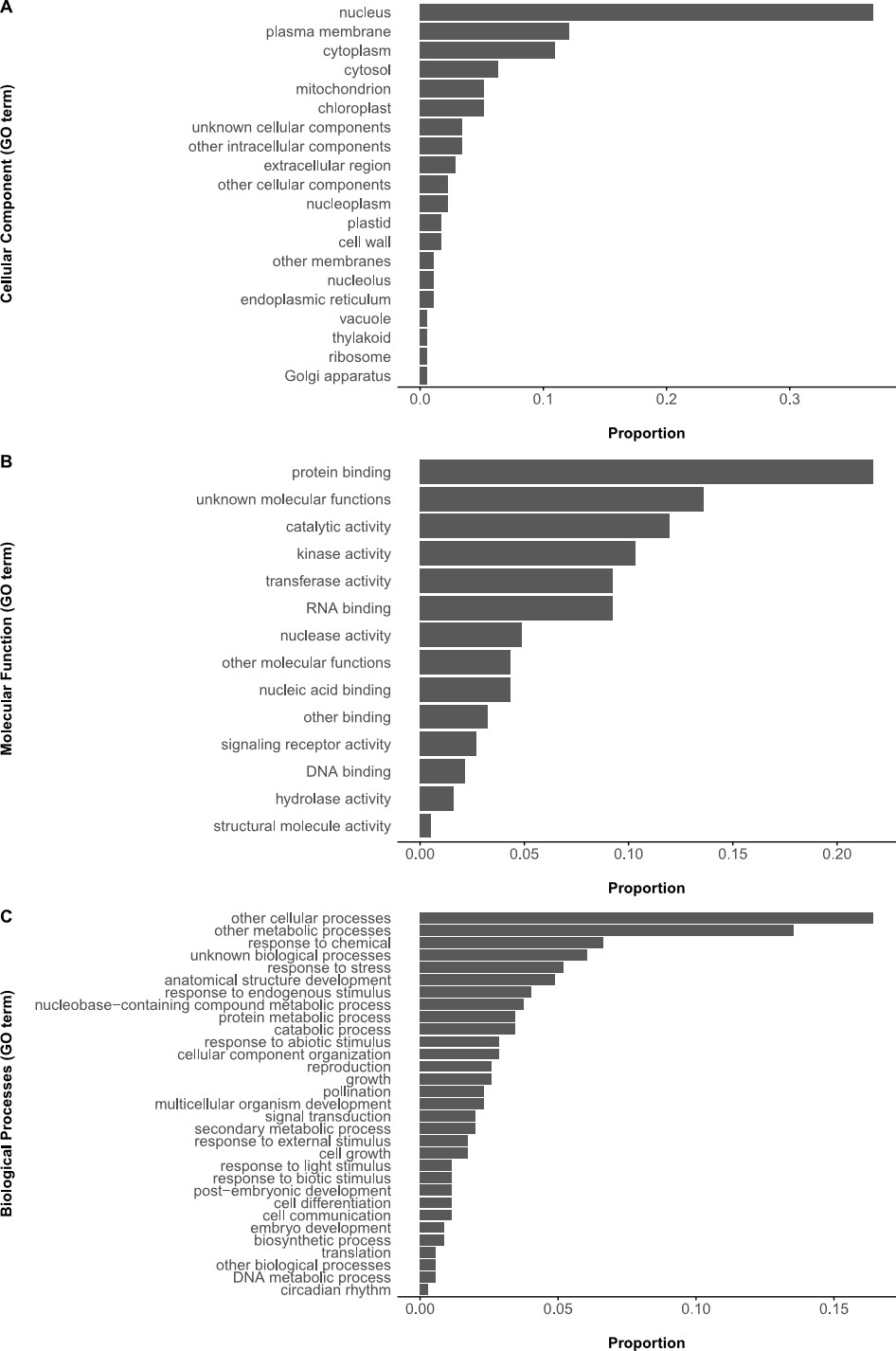

### Appendix S9. GO annotations enrichment and depletion analysis. Listed are GO categories, GO terms, log-transformed odd ratios (Log OR), Haldane-Anscombe-corrected log-transformed odd ratios (HA-corrected Log OR), and estimated FDRs.

| **GO category*** | **GO term** | **Log OR**** | **HA-corrected Log OR** | **FDR** |
| --- | --- | --- | --- | --- |
| CC | plasma membrane | 1.72 | 1.70 | <0.001 |
| MF | signaling receptor activity | *Inf* | 3.26 | 0.053 |
|  | DNA-binding transcription factor activity | *-Inf* | -2.63 | 0.084 |
| BP | biosynthetic process | -1.90 | -1.76 | 0.001 |
|  | cell growth | *Inf* | 3.64 | 0.002 |
|  | growth | 2.58 | 2.41 | 0.001 |
|  | pollination | 3.16 | 2.81 | 0.001 |
|  | protein metabolic process | 1.26 | 1.25 | 0.030 |
|  | cell cycle | *-Inf* | -3.15 | 0.001 |
|  | lipid metabolic process | *-Inf* | -2.57 | 0.032 |

* CC: Cellular Component, MF: Molecular Function, and BP: Biological Process.

** *Inf* and *-Inf* results from odd ratios calculated from 0 success probabilities accounted for with Haldane-Anscombe correction.

### Appendix S10. Distribution of genotype frequencies (0/0, 0/1, 1/1) at the shared locus (locus_4218 [APFE030529380.1:392010]) for each Torrey pine ancestry (Mainland [75 individuals], Island [68 individuals], F1 hybrid [66 individuals]).

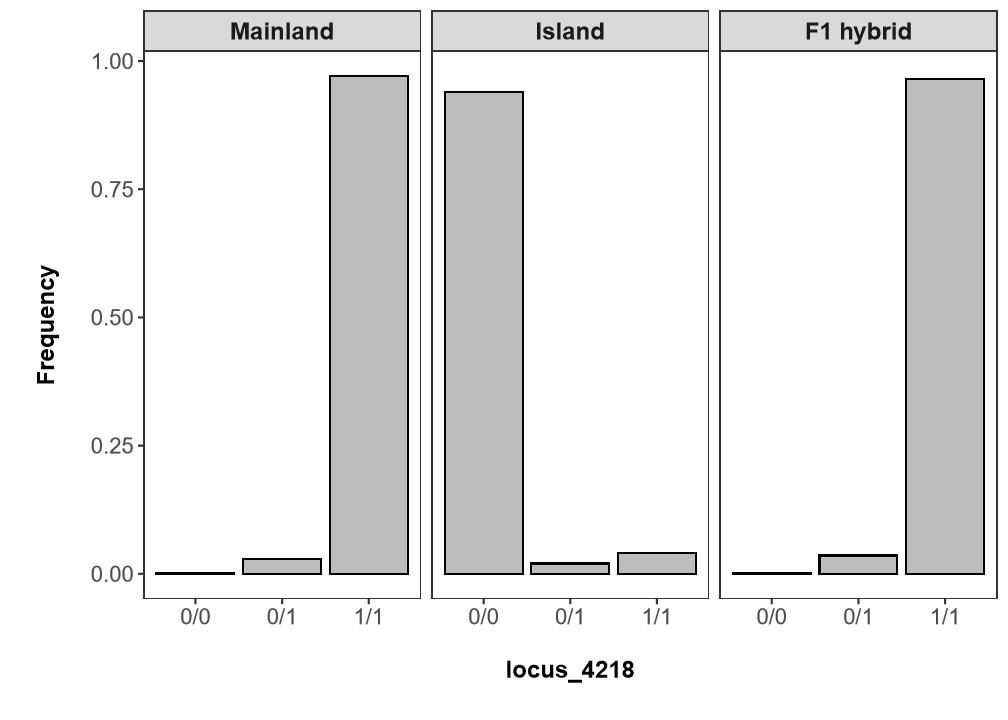

### Appendix S11. Summary statistics associated with evaluating statistical differences in expected marginal means of measured phenotypes between genotypes. Listed are the phenotypic trait assessed (Phenotypic trait), the pair of genotypes considered (Contrast), the difference in expected marginal means observed between genotypes (EMM difference), as well as the degree of freedom (df), test statistic (T ratio), and adjusted P value (Adj. P value) associated with every phenotype-specific pairwise two-sample unpaired t-test performed.

| **Phenotypic trait** | **Contrast** | **EMM difference** | **df** | **T ratio** | **Adj. P value*** |
| --- | --- | --- | --- | --- | --- |
| Tree height (cm) | 0/0 - 0/1 | -45.90 | 166 | -1.797 | 0.174 |
|  | 0/0 - 1/1 | -59.70 | 165 | -4.065 | <0.001 |
|  | 0/1 - 1/1 | -13.80 | 166 | -0.596 | 0.823 |
| Number of conelets | 0/0 - 0/1 | -2.16 | 166 | -2.224 | 0.070 |
|  | 0/0 - 1/1 | -3.55 | 165 | -6.341 | <0.001 |
|  | 0/1 - 1/1 | -1.39 | 166 | -1.569 | 0.262 |
| Number of immature cones | 0/0 - 0/1 | -0.64 | 166 | -2.991 | 0.009 |
|  | 0/0 - 1/1 | -0.88 | 165 | -7.139 | <0.001 |
|  | 0/1 - 1/1 | -0.24 | 166 | -1.232 | 0.436 |
| Number of mature cones | 0/0 - 0/1 | -0.60 | 166 | -2.114 | 0.090 |
|  | 0/0 - 1/1 | -1.13 | 165 | -6.914 | <0.001 |
|  | 0/1 - 1/1 | -0.53 | 166 | -2.054 | 0.103 |

* P values are adjusted internally within the function *emmeans()* using the Tukey method for comparing a family of three estimates.

### Appendix S12. Summary statistics associated with evaluating statistical differences (A) in the degree of fixation estimated from the whole genomic data set (All) among Torrey pine ancestries (F1 hybrid, island, mainland), and (B) in observed heterozygosity (H_O_) averages among Torrey pine ancestries within SNP sets (All, GWAS, Low).

**A**. Summary statistics associated with Fisher’s exact tests for count data. Listed are the dimensions of contingency tables used (Dimension), ancestry pairs used for pairwise comparisons when using 2 by 2 contingency tables (Contrast), estimated odd ratios and their 95% confidence intervals (Odd ratio), and significance parameters (p-value when using the 3 by 2 contingency table, FDR otherwise).

| **Dimension** | **Contrast** | **Odd ratio**  **(95% CI)** | **P value / FDR** |
| --- | --- | --- | --- |
| 3 by 2 | - | - | < 0.001 |
| 2 by 2 | Island - F1 hybrid | 4.34  (3.65 - 5.18) | < 0.001 |
|  | Mainland - F1 hybrid | 4.03  (3.38 - 4.81) | < 0.001 |
|  | Mainland - Island | 0.93  (0.83 - 1.04) | 0.19 |

**B**. Summary statistics associated with Dunn’s tests. Listed are the genomic data sets used for analysis (SNP set), ancestry pairs used for pairwise comparisons (Contrast), test statistics (Z-statistic), and false discovery rates (FDR).

| **SNP set** | **Contrast** | **Z-statistic** | **FDR** |
| --- | --- | --- | --- |
| All | F1 hybrid - Island | 0.46 | 1 |
|  | F1 hybrid - Mainland | 0.15 | 0.88 |
|  | Island - Mainland | -0.31 | 1 |
| GWAS | F1 hybrid - Island | 2.33 | 0.06 |
|  | F1 hybrid - Mainland | 1.78 | 0.11 |
|  | Island - Mainland | -0.55 | 0.58 |
| Low | F1 hybrid - Island | -5.38 | < 0.001 |
|  | F1 hybrid - Mainland | -5.62 | < 0.001 |
|  | Island - Mainland | -0.24 | 0.81 |

### Appendix S13. Distribution of observed heterozygosity estimated at 11,379 SNPs across 209 Torrey pine samples, including 66 F1 hybrid (brown), 68 island (yellow), and 75 mainland (orange) individuals.

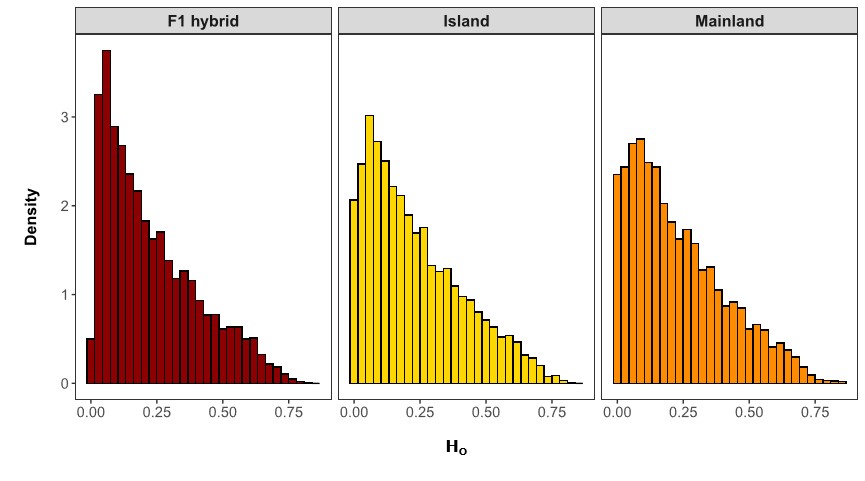

### Appendix S14. Distribution of locus-specific island-mainland pairwise Nei’s F_ST_ values* for two different SNP sets (All, Low). All: data set containing all retained SNPs after filtering (11,379 SNPs), Low; data sets containing SNPs exhibiting reduced heterozygosity relative to expectations (185 SNPs). Sample size is 143 individuals (68 island and 75 mainland individuals) for both SNP sets.

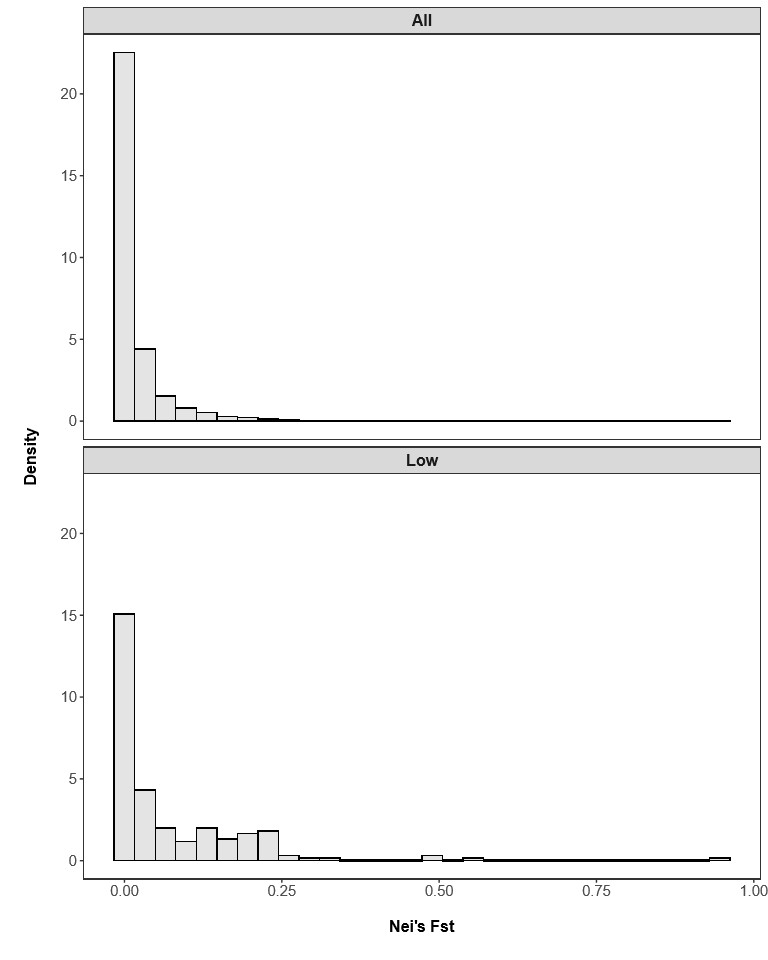

*Pairwise Nei’s F_ST_ values were estimated in R using the function *basic.stats()* implemented within the package *hierfstat*.
